# The Microbiome and Volatile Organic Compounds Reflect the State of Decomposition in an Indoor Environment

**DOI:** 10.1101/2022.05.18.492585

**Authors:** Veronica M. Cappas, Emily R. Davenport, Dan G. Sykes

**Affiliations:** Forensic Science Program, Eberly College of Science, The Pennsylvania State University, State College, PA; Department of Biology, Huck Institute of Life Sciences, The Pennsylvania State University, State College, PA; Department of Chemistry, Eberly College of Science, The Pennsylvania State University, State College, PA

## Abstract

Because of the variety of factors that can affect the decomposition process, it can be difficult to determine the post-mortem interval (PMI). The process is highly dependent on microbial activity, and volatile organic compounds (VOCs) are a by-product of this activity. Given both have been proposed to assist in PMI determination, a deeper understanding of this relationship is needed. The current study investigates the temporal evolution of the microbiome and VOC profile of a decomposing human analog (swine) in a controlled, indoor environment. Microbial communities and VOCs were sampled at six-time points, up to the active decay phase. Sampling locations included the abdominal area, anus, right ear canal, and right nostril. Bacterial communities were found to significantly change during decomposition (p-value < 0.001), and communities evolved differently based on sampling location. The families Moraxellaceae, Planococcaceae, Lactobacillaceae, and Staphylococcaceae drove these community shifts. From random forest analysis, the nostril sampling location was determined to be the best location to predict stage of decomposition. Individual VOCs exhibited large temporal shifts through decomposition stage in contrast to smaller shifts when evaluated based on functional groups. Finally, pairwise linear regression models between abdominal area bacteria and selected VOCs were assessed; Planococcaceae and Tissierellaceae were significantly correlated to indole. Overall, this study provides an exploratory analysis to support the connection between the microbiome, VOCs, and their relationship throughout decomposition.

**Importance:** This research provides valuable insight into the complex process of decomposition, which is pertinent to forensic death investigations. The temporal evolution of both the microbiome and volatile organic compounds (VOCs) were characterized as a function of stage of decomposition and evaluated their interdependency upon one another. In turn, this information may assist in determining time since death, and fill a knowledge gap about VOC-bacteria associations during the decay process.

## I. Introduction

Determining time since death or the post-mortem interval (PMI) is an important aspect to forensic death investigations in order to help build a timeline of the event in question. Early time since death indicators include algor mortis (stabilization of body temperature with environment), rigor mortis (stiffening due to the lack of ATP production), and livor mortis (settling of blood) (1). Later physical signs of decomposition like tissue state, gas build-up (bloating), fluid purging, insect colonization, botanical growth, and skeletal chemical changes have also been used to determine PMI (1, 2). Because decomposition is highly variable, it can be difficult to determine time since death, and common determinations of PMI can have limitations (1). Algor mortis can be affected by decomposition-induced (or environmental) temperature changes, insect activity can be limited in circumstances like indoors or certain climates, and rigor mortis disappears after 36 hours (1, 3). To overcome some of these limitations, the use of microorganisms along with volatile organic compounds have been proposed for PMI determination (2, 4).

Generally, decomposition begins in high-enzyme content organs due to autolysis, which is the break-down of tissue and release of cellular contents due to digestion from the body’s own enzymes. Next, putrefaction occurs from bacteria, other microorganisms, and macroorganisms consuming the body’s nutrients (5). Indigenous bacteria present in places like the gastrointestinal tract and the respiratory system begin to flourish after the immune system fails and can then migrate to other tissues (6, 7); bacteria from the environment can colonize the host and become a part of the decomposition community. As the microbial action breaks down tissue into simpler compounds, new and additional food resources are created which promote further colonization and growth creating a complex changing community (7, 8).

This decomposition process creates a unique environment that depends on bacterial interactions, density of communities, and the availability of nutrients, which can be affected by location on the body and where the body itself is located (9). Because there can be significant shifts in microbial communities due to resource availability and interactions, such shifts have been supported to correlate with time since death (10-20). For example, there are often shifts in communities associated with the bursting event, which marks the transition between the bloat and post-bloat stages (10, 12, 16).

Gases that induce bloating and that are responsible for the odor of decomposition are classified as volatile organic compounds (VOCs) (21). VOCs are, in part, a by-product of microbial action, and the temporal evolution of the VOC profile may be an important marker for PMI indication (22-24). As such, there is an extensive database correlating VOC profiles with different environmental conditions including body location (burial versus surface placement), physicochemical conditions (temperature, season, humidity, etc.), along with stage of decomposition (4, 23, 25-32).

In contrast, we lack an understanding of the connection between shifting microbial communities and VOCs in the context of decomposition. The database, “m.VOC,” served as a warehouse for information on the VOC emissions from specific bacterial communities. However, the database is limited in scope, currently inaccessible, and does not necessary reflect VOCs gathered from the decomposition environment (33). A study measured the VOC production of three bacteria associated with different stages of decomposition over a five-day period, showing a high-degree of specificity between VOC production and bacterium (21). The study demonstrates the utility of examining VOCs in relation to the decomposition bacteria; however, the understanding of the VOCs produced by the majority of microbes involved in decomposition needs further exploration (21). In addition, specific VOCs and bacteria families were identified and significantly correlated through stages of decomposition of still-born piglets (34). At present date, these few studies are the only research attempts to directly link decomposition, microorganisms, and VOCs.

Furthermore, few studies directly focus on bacterial succession or the temporal change in VOC profiles during decomposition in an indoor environment. An indoor environment is common, if not more common than an outdoor environment for death scenes, and, therefore, warrants further investigation (35). Microbiome response and VOC production can be studied without fluctuating variables that affect decomposition, like temperature, weather, and insect activity. This can better pinpoint direct effects from bacteria on decomposition and determine the extent to which external variables impact the decomposition microbiome.

Here, these gaps are addressed by exploring the microbiome and VOC profile in parallel of a human analog (swine) in a controlled, indoor environment. We find that both the microbiome and VOC profile transform throughout decomposition, while also discovering potential associations between bacteria families and selected VOCs. Results could further help deconvolute the complex process of decomposition and support the use of the microbiome and VOCs to determine the post-mortem interval.

## II. Materials and Methods

### A. Swine Placement and Experiment Set-Up

A male Yorkshire swine, obtained from The Pennsylvania State University’s (PSU) Swine Facility, was euthanized via captive bolt. The swine was approximately 35 pounds and 8 weeks old. PSU’s Institutional Animal Care and Use Committee (IACUC) (approval ID: PROTO202001382) reviewed and approved the study. Swine are generally accepted as human analogs for decomposition studies because of their similar body composition and anatomy, similar intestinal microbiome, and similar decay process to humans (36, 37). In addition, human cadaver availability can be limited. Immediately after euthanization, the swine was placed in an indoor enclosure (**Fig. S1**). The enclosure was constructed of plexiglass and fitted with a two-way pass-through box, gloves, and port for VOC sampling. The pass-through box and gloves ensured minimal outside contamination to the swine. The enclosure was placed in a chemical fume hood. Once the swine was placed in the enclosure, it remained sealed during the decomposition process. The temperature throughout the study was maintained at approximately 73 degrees Fahrenheit (°F).

The microbiome of and VOCs emitted by the swine were sampled at six time points (**Fig. 1**). The stage of decomposition was approximated using the total body scoring system from Megyesi et al. 2015 (38).

**Figure 1:**
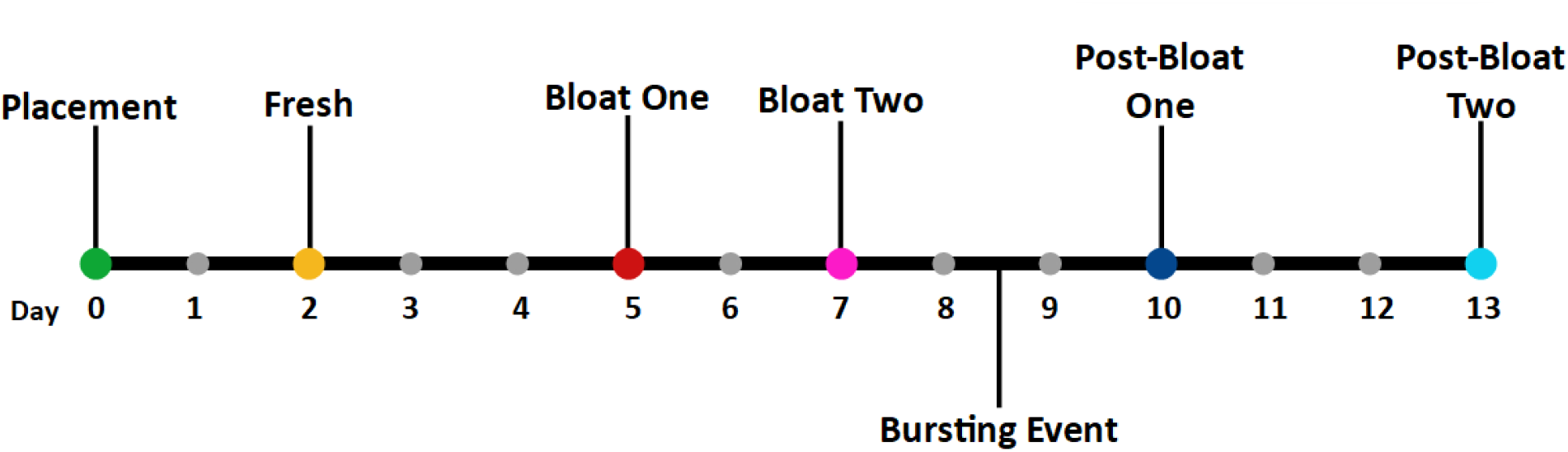
Timeline with sampling days (colored dots) and corresponding sampling day labels. The bursting event (between days 8 and 9) is the rupturing around the abdominal area from the build-up of gases; internal organs became exposed.

### B. Microbiome Sampling

For every sample day, four locations were sampled in triplicate: abdominal area, anus, right ear canal, and right nostril. These locations were selected to minimize any disturbance to the swine during sampling. This is an important aspect when sampling during investigations, as minimal alternation to evidence is high priority. Nylon flock swabs (BD; Franklin Lakes, NJ) were used to sample the microbiome. Before swabbing, the swab was dipped in sterile, phosphate buffer solution. After swab collection, each tip was removed with sterile scissors and place in a sterile, plastic collection tube. Control swabs were collected from the ambient air inside of the chamber by leaving a swab exposed for at least 20 seconds. The samples were stored at -20 degrees Celsius (°C) until DNA extraction was completed. Due to the bursting event, the skin ruptured and the internal organs were exposed. After the bursting event, exposed internal organs were swabbed in addition to the abdominal area skin for the abdominal area sampling location.

### C. Microbial DNA Extraction, PCR, and Sequencing

The UltraClean Microbial DNA Kit (Qiagen; Germantown, MD) was used to extract DNA following a modified version of the manufacturers recommended protocol. Swabs were vortexed with PowerBead solution, and the swabs were removed from the collection tubes before proceeding with the recommended protocol. In addition, low TE buffer was used instead of the Solution EB for the final elution step. Extracted DNA was quantified using the Qubit 1X dsDNA HS Assay Kit and Qubit Quantification system (Qiagen; Germantown, MD). The ZymoBIOMICS Microbial Community Standard (Zymo Research; Irvine, CA) was used as a positive extraction control, and a negative extraction control was also performed. The V4 region of the 16S ribosomal ribonucleic acid (rRNA) gene was amplified from the extracted DNA, as it is a commonly used phylogenetic marker gene for bacteria and archaea. This region of the 16S gene is hypervariable between species, which qualifies as a good region for identification (14, 17). Primers used include 515F-v2 (Parada) (5′ -TCGTCGGCAGCGTCAGATGTGTATAAGA GACAGGTGYCAGCMGCCG CGGTAA -3′) and primer 806R-v2 (Apprill) (5′-GTCTCGTGG GCTCGGAGATGTGTATAAGAGACAGGGACTACNVGGGTWTCTAAT -3′) (Integrated DNA Technologies; Coralville, Iowa). These primers contained Illumina overhangs. Samples were prepared for PCR by combining 2 microliters (µL) of extracted sample DNA (2.5 ng/ µL), 0.8 µL of 10 µM paired primer stock, 10 µL of 2x Invtirogen Platinum SuperFi Master Mix (ThermoFisher Scientific; Waltham, MA), and 7.5 µL of sterile PCR water into a 20-µL PCR reaction tube. If a DNA extracted sample was below 2.5 ng/ µL, water input was reduced in the reaction. Samples were heated for 2 minutes at 99 °C followed by 25 cycles of 10 seconds at 98 °C, 56.5 °C for 20 seconds, and 72°C for 15 seconds. Finally, a 5-minute hold at 72°C (39). The ZymoBIOMICS Microbial Community DNA Standard (Zymo Research; Irvine, CA) was used as a positive PCR control, along with a negative PCR control.

Amplification was verified via gel electrophoresis by comparing randomly selected samples to a reference DNA ladder at about 250 base pairs. Samples were sent to the Huck Institutes of the Life Sciences’ Genomics Core Facility (The Pennsylvania State University, University Park, PA) for sequencing. The Genomics Core Facility followed a modified protocol of the “16S Metagenomic Sequencing Library Preparation” and performed a second round PCR where the rest of the Illumina adapters and indexes were added (40). After the second round of PCR, the concentration of the libraries was normalized using Mag-Bind® Equipure Library Normalization Beads (Omega Bio-Tek; Norcross, GA). An approximate equimolar pool of the libraries was made by pooling an equal volume of each library. The pool was sequenced using 250 × 250 paired-end sequencing run on an Illumina MiSeq (Illumina; San Diego, CA).

### D. Microbiome Bioinformatics Analysis

A total of 19.8 million 251 base pair paired-end reads were generated, with the average read depth of 106,000 per sample. The raw sequenced reads were processed by PSU’s Bioinformatics and Genomics Consulting Center. Adapter sequences and low-quality bases were trimmed using the paired-end mode in Trimmomatic (USADEL Lab; version 0.39) (41). Low-quality bases were cut that fell below Q20 within a 4 base pair window, based on a sliding window approach. In addition, reads less than 35 base pairs were filtered after trimming. 18.6 million reads with an average read depth of 100,000 per sample were retained after trimming and filtering. Trimmed reads were run classified with Kraken2-Bracken (The Center for Computational Biology at John Hopkins University; Baltimore, MD; version 2.1.1) using the Greengenes database (Lawrence Berkeley National Laboratory (LBNL); Berkeley CA; version gg_13_5) (42, 43). Raw sequence reads are stored at the National Center for Biotechnology Information (NCBI) Sequence Read Archive (SRA) (BioProject: PRJNA827913). Data statistics before and after trimming, FastQC reports before and after trimming, MultiQC reports before and after trimming, commands for the Kraken2-Bracken analysis, and classification reports are stored on the Zenodo data repository. For downstream analysis, the genus classification output from the Greengenes database (LBNL; Berkeley CA; version gg_13_5) was used and uploaded to R (43, 44). This output includes the operational taxonomic unit (OTU) matrix, taxonomy table, and metadata and is located in **File S1**.

The “phyloseq” package in R was used for analysis of classification output (44, 45). OTUs with an abundance of less than 0.001% if total read count were filtered. In addition, sampled were rarefied down to 403 OTUs from 1,181 OTUs and 50,476 reads per sample (seed 333). Samples Ln67 and EBec33 were not uploaded into R and removed from the dataset because of their extremely initial low read count (< 5 reads). Alpha diversity was assessed using the Shannon Diversity Index using the “phyloseq” package (45). Both taxa richness, the number of taxa, and taxa evenness, how the abundance of taxa are distributed, were used to evaluate the diversity of communities within a sample. Each sampling location was assessed over the sampling period, and each sampling location was compared at each decomposition stage. Alpha diversity box plots were also produced using the “phyloseq” package (45). Diversity indices were compared as previously described using the Kruskal-Wallis Rank Sum Test and Dunn’s Kruskal-Wallis Multiple Comparisons using the “stats” and “FSA” packages (44, 46). Significance was set at 0.05 for adjusted p-values, using the Benjamini-Hochberg method.

The diversity of communities between samples, known as beta diversity, was evaluated in three ways. First, using Bray-Curtis dissimilarity, a Principal Coordinate Analysis (PCoA) plot was produced using the “phyloseq” and “ggplot2” packages (45, 47). Using Bray-Curtis Dissimilarity and permutational multivariate analysis of variance (PERMANOVA), beta diversities were compared by both decomposition stage and sampling location using the “vegan” package and “adonis” function (48). The second and third approaches included relative abundance bar charts of taxa against stage of decomposition and line graphs of top changing taxa which were produced using “ggplot2” (47). Machine learning using random forest for each location was performed using the “randomForest” package (49, 50). Four different groupings of decomposition stages were tested as response variables. Grouping 1 was each original decomposition stage. Grouping 2 separated the Placement and Fresh stages and merged the Bloat and Post-Bloat stages. Grouping 3 merged all the Pre-Bloat (Placement and Fresh), Bloat, and Post-Bloat stages together. Finally, Grouping 4 was based on stages grouped before and after the bursting event.

### E. VOC Sampling and Gas Chromatography-Mass Spectrometry

The indoor enclosure head space was sampled using, a 65-µm DVB/ PDMS Solid Phase Microextraction (SPME) fiber (Restek; Bellefonte, PA). Using the port at the top of the enclosure, the SPME fiber was inserted into the headspace of the enclosure and was then processed using gas chromatography-mass spectrometry (GC-MS) (4). Again, the enclosure headspace was sampled at the same six timepoints as the microbiome. The enclosure headspace was sampled before swine placement for a blank or control sample to ensure accurate deconvolution of the VOCs emitted from the swine from ambient VOCs in the laboratory air. The capillary column was a 30-m L×0.25-mm ID×0.25-µm film thickness RTx-5ms (Restek; Bellefonte, PA). SPME fibers were inserted into a heated injection port (260°C), and a splitless injection method was employed using a 2-minute hold time. The oven temperature gradient was employed: initial temperature of 35°C held for 2 minutes then ramped at 15°C/min to 260°C, which was held for 10 minutes. The MS was scanned at mass range of 35-450 m/z at 200°C. SPME fibers were cleaned by inserting into the injection port for 30 minutes at 260°C.

### F. VOC Data Analysis

Chromatographic peaks were identified using National Institute of Standards and Technology (NIST) Mass Spectral libraries 21 and 107. A peak was confirmed if the match threshold was at least 70. If a compound from a sample day was seen in the blank collection, it was not considered to be a compound of interest and excluded from **Table S1**. Retention time, peak area, and logarithmic values of relative peak areas for each VOC from each stage are stored in **File S2**. VOCs were also categorized into functional groups. Principal component analysis (PCA) plots were created using Minitab Statistical Software (Minitab, LLC; State College, PA). For the PCA plots, relative peak area was determined for each compound by dividing peak area by sum of peak areas from a sample, followed by a logarithmic transformation (51). The absolute values of relative peak areas were then used for the PCA plots. These relative peak areas were determined only for general comparison of VOC profile composition between stages and not for any quantitative conclusions.

### G. Correlation Analysis between the Microbiome and VOC Profile Analysis

To investigate correlation between trends of the microbiome and the VOC profile, linear regression was employed. Simple linear regression models were fitted in a pairwise manner with each selected VOC attempting to be explained by a bacteria taxon. The top 30 most prevalent families were selected from the abdominal area sampling location (**Table S3**). The selected VOCs from the overall swine profile included nonanal, indole, dimethyl disulfide, dimethyl trisulfide, phenol, butanoic acid, and pentanoic acid. Preliminary data analysis and linear regression was performed using R with the “stats,” “base,” and “graphics” packages (44). First, clusters of selected VOCs were created based on scaled trends of VOCs. In addition, correlations, based on Pearson’s correlation coefficient, among selected VOCs and prevalent abdominal area genera were produced individually. The VOC correlation plot was produced using the “corrplot” package (52). P-values from the bacteria-VOC pairwise linear regression models were extracted to assess if correlations were significant. However, due high amount of pairwise correlations performed, p-values were adjusted using the Holm method to correct for any significant correlations found by chance, as known as Family-Wide Error Rate (53). Significance was defined as a p-value of 0.05 or lower. Finally, the “pheatmap” package created visual graphics of correlation trends between selected VOCs and the prevalent bacteria families and genera (54). Clusters were based on Pearson’s correlation coefficient.

### H. Data Availability

Raw sequence reads stored at NCBI’s SRA (BioProject: PRJNA827913) can be accessed at *https://dataview.ncbi.nlm.nih.gov/object/PRJNA827913?reviewer=9c4vl7q2jm72hcbpul2b1hscj8(reviewerlink)*. Pre-analysis bioinformatics files as described in the **Microbiome Bioinformatics Analysis** sub-section are also located at the Zenodo data repository (DIO: 10.5281/zenodo.6370443). Supplemental Files including **File S1, S2**, and **S3** are located on the Zenodo data repository (DIO: 10.5281/zenodo.6539755).

## III. Results

### A. Microbiome

#### i. Assessing controls reveals effective experimental design

The positive extraction and PCR positive samples performed within specifications of the manufacture. Community composition of air control samples are relatively clustered together and are located near some experimental samples, indicating composition similarity **(Fig. S2**). However, this may not be surprising because of exchange between the air bacteria and bacteria originating from the swine. The air control samples and the extraction negative control samples that had higher reads than expected may be explained by the normalization process during library preparation (**File S1**). For the control analysis, taxa information uploaded into R was not filtered or rarefied. In addition, quantification values after extraction were “too low” to read from the Qubit Quantification system (Qiagen; Germantown, MD), further supporting the high read count for controls occurred because normalization step and not contamination.

#### ii. Sample taxa diversity does not reveal definitive correlation with stage of decomposition

Alpha diversity was assessed through the Shannon Diversity Index. Each sampling location was independently measured through decomposition stages (**Fig. 2**). Few definitive conclusions can be made on individual sample diversities in an indoor environment throughout decomposition (**Fig. 2** and **Table S2**). Out of all the locations, the abdominal area had the most definitive trend with community diversities dropping in the Post-Bloat stages (**Fig. 2a)**. In addition, the ear canal’s alpha diversity dropped after the Placement stage; however, it was interesting that diversity then again increased after the Bloat One stage. Due to overlapping ranges, patterns were difficult to conclude for the anus and nostril sampling locations. All sampling locations except the anus’s alpha diversities were significantly different by decomposition stage (p-value < 0.05) (**Table S2**). However, any significant changes between specific stages from the Dunn’s Multiple Comparison Test were not common and only observed between stages before and after the bursting event (**Fig. 3** and **Table S2**).

**Figure 2:**
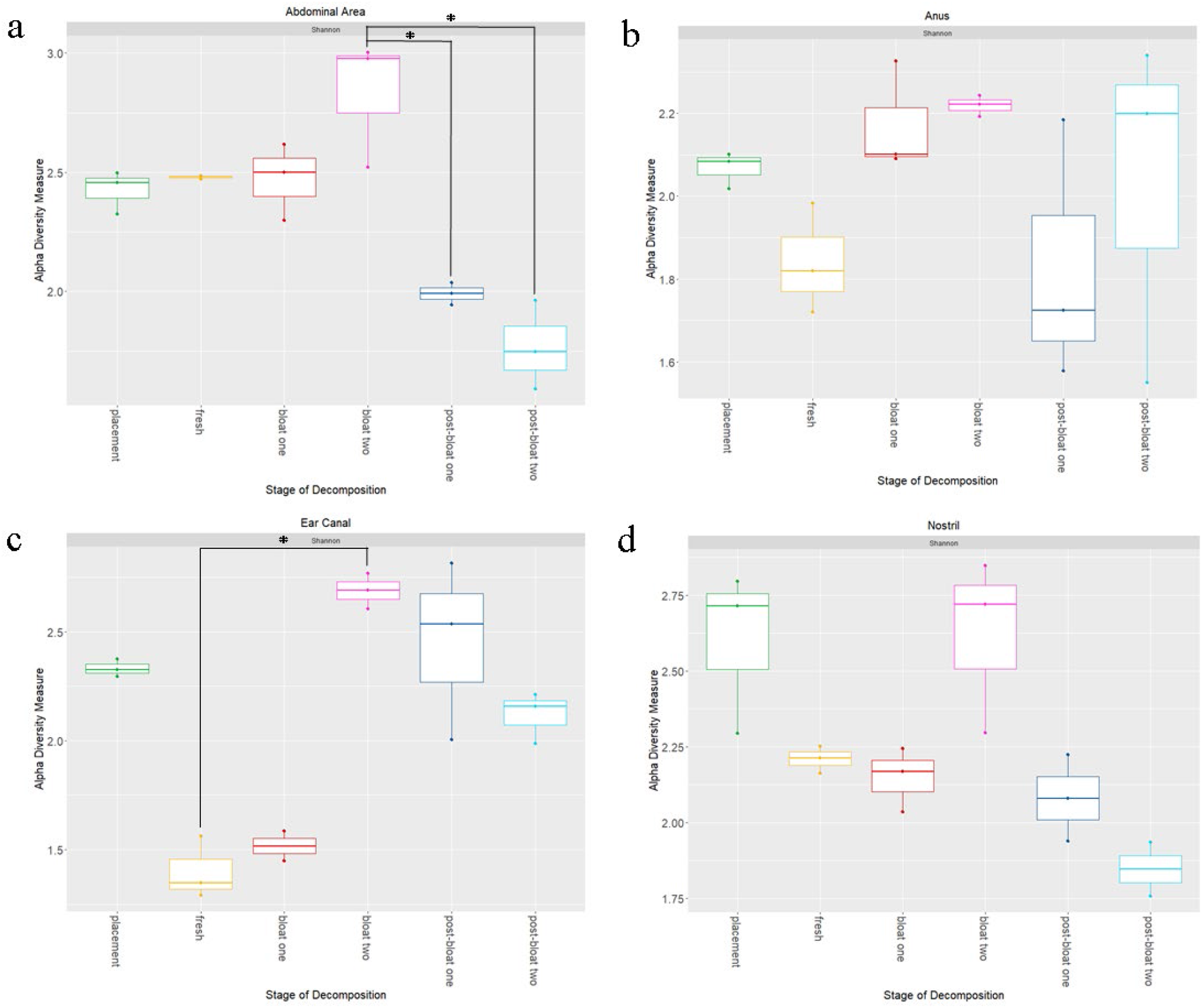
Alpha Diversity Box Plots of Shannon Diversity Index Measurements versus Stage of Decomposition for **a**. Abdominal Area, **b**. Anus, **c**. Ear Canal, and **d**. Nostril. Higher indices indicate higher taxa richness and evenness in taxa abundance. Points on boxes indicate specific sample measurements. Significant stage differences based on Dunn’s Multiple Comparison Test (p < 0.05) are denoted by “*.”

**Figure 3:**
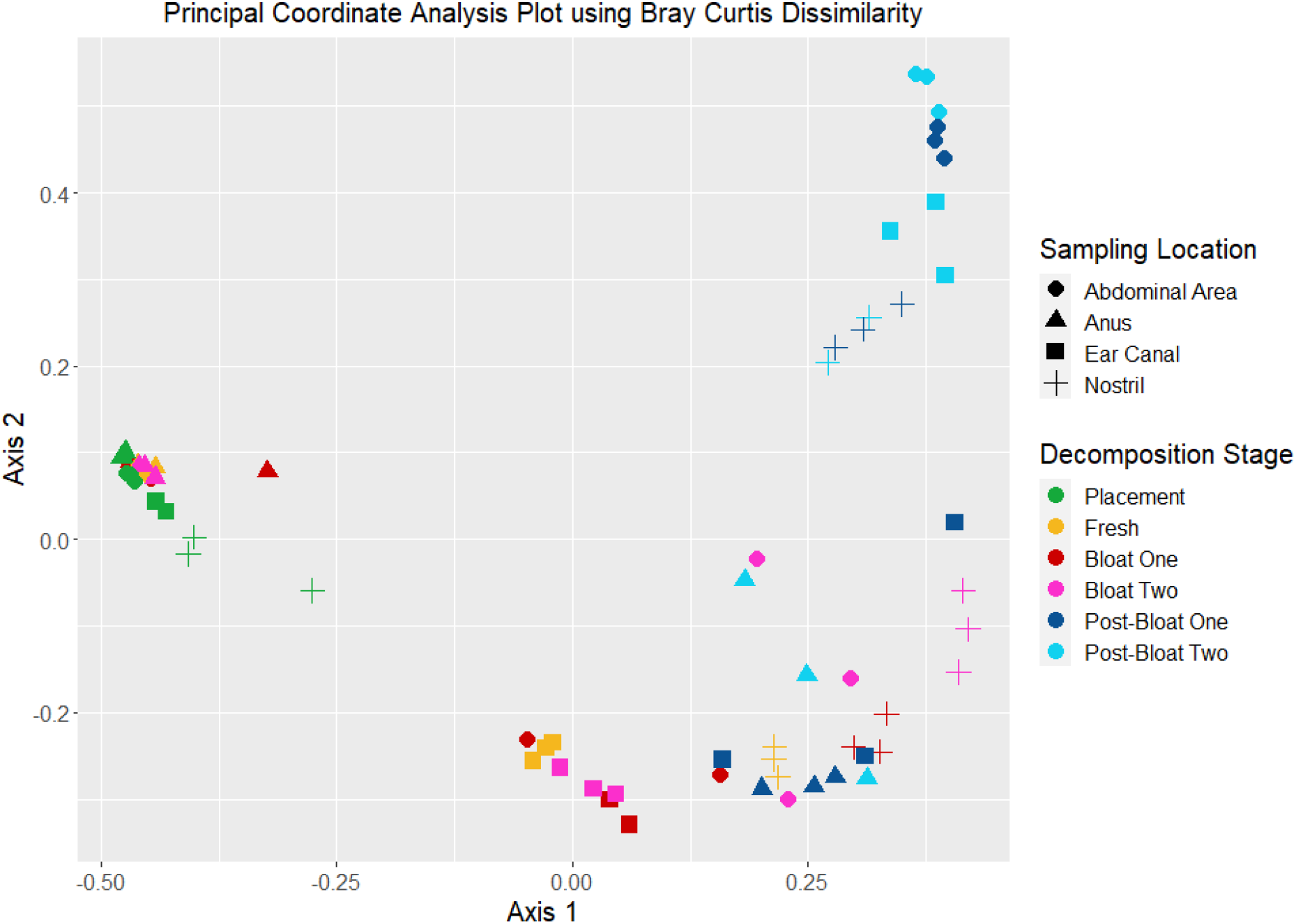
PCoA plot of experimental microbiome samples produced from the first two principal coordinate eigenvector values. Dissimilarity (or similarity) of sample communities through spatial distances from Bray-Curtis Dissimilarity at the genus level between samples are shown. Swab/ sampling location is denoted by shape, and decomposition stage is denoted by color.

Comparison of swab locations for each stage showed clearer trends (**Fig. S3**). In earlier stages, the abdominal area and nostril locations had higher alpha diversities, while in later stages the ear canal had higher alpha diversity measurements (**Fig. S3**). However, only the Fresh stage had significantly different alpha diversity based on sample location (p-value < 0.05). The abdominal area and the ear canal in the Fresh and Bloat One stages were significantly different from the Dunn’s Multiple Comparison Test (p-value < 0.05) (**Table S3**).

#### iii. Bacterial communities show significant composition differences between stage of decomposition and sampling locations

Beta diversity, in contrast to alpha diversity, compares bacterial community composition between samples. To assess differences in community composition, PERMANOVA results of both stage of decomposition and location were performed. Community composition was significantly different between decomposition stages and sampling locations (Stage of Decomposition: Sum of Squares=7.3396, F-statistic=45.967, p-value < 0.001; Sampling Location: Sum of Squares=4.9199, F-statistic=51.355, p-value < 0.001; Stage of Decomposition and Sampling Location: Sum of Squares=6.8747, F-statistic=14.352, p-value < 0.001). This is concordant with the clusters observed in the PCoA plot (**Fig. 3**) and visually different community compositions seen in the relative abundances (**Fig. 4**), depicting a change in communities through decomposition. Triplicate samples were relatively clustered, indicating that the replicates had minimal variation (**Fig. 3**). Despite sampling location, all Placement samples plot near one another. In addition, some sampling location Fresh and Bloat samples also clustered near the Placement samples. Pre-Bursting ear canal sampling locations tended to cluster distinctly from other locations. Several sampling locations’ Post-Bloating samples were clustered near one another. Post-Bloat samples for the anus, Post-Bloat One stages for the ear canal, Fresh and Bloat samples for the nostril, and Bloat samples for the abdominal area clustered loosely, but not as tight as the previously identified clusters.

**Figure 4:**
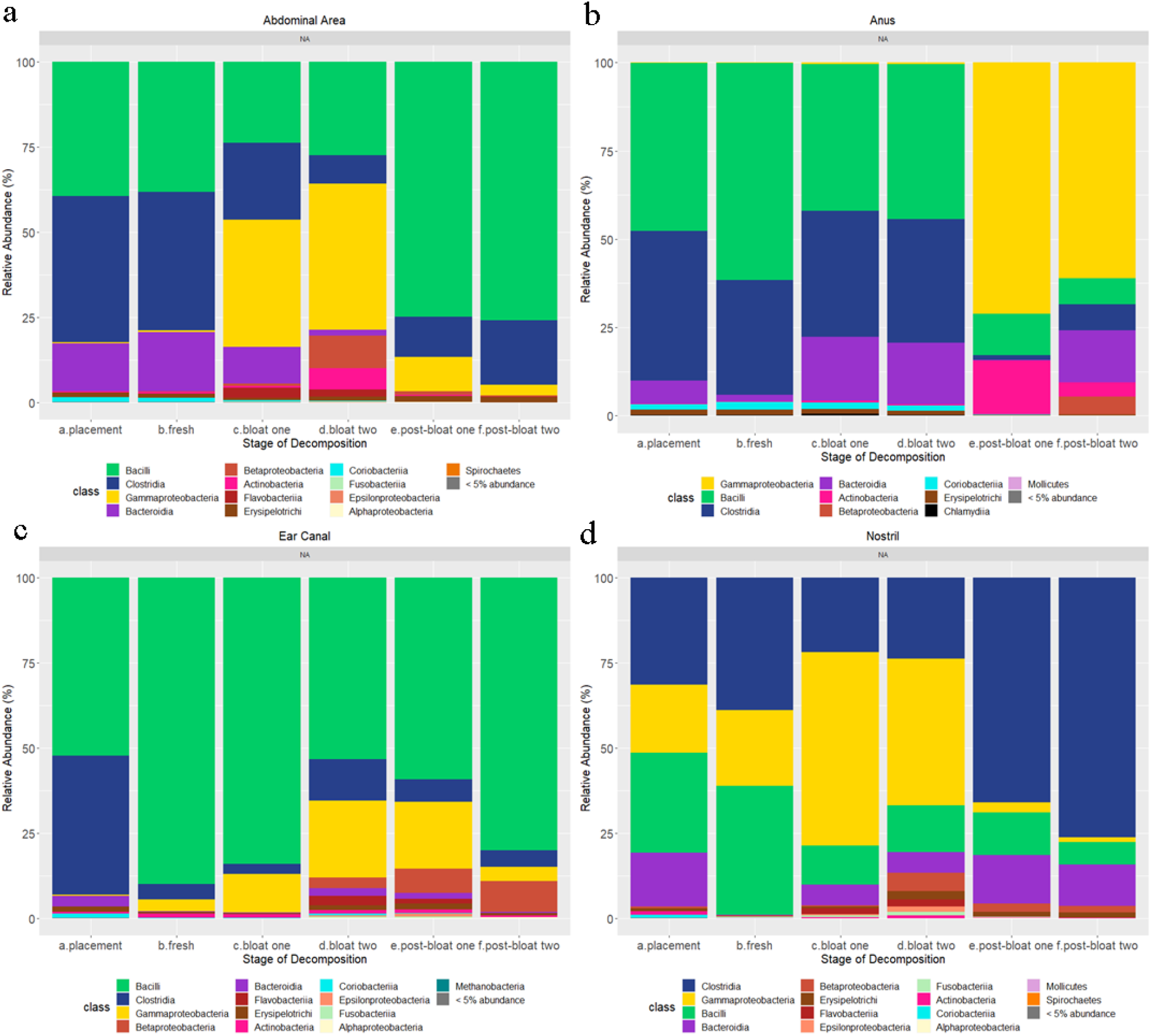
Relative Abundance (%) of Classes of Each Stage of Decomposition for **a**. Abdominal Area, **b**. Anus, **c**. Ear Canal, and **d**. Nostril Relative sampling locations. Relative abundance was defined as the amount of reads for a class divided by the total amount of reads in a stage of decomposition. Classes that represented less than 5% of the population of identified taxa for a stage were merged into one class category.

The drivers or determinants of these clusters can be attributed to abundances of prevalent taxa present in the samples (**Fig. 4** and **5**). Gammaproteobacteria, a diverse class, Bacilli, a class consisting of the Bacillales and Lactobacillales orders, and Clostridia, a mostly anaerobic class, were the most prevalent classes regardless of the sampling location. Minor classes include Bacteriodia, Betaproteobacteria, and Actinobacteria, Flavobacteria, and Erysipelotrichi. Many families identified in **Fig. 5** are a part of the major classes in **Fig. 4** and coincide with community shifts. For example, in the abdominal area, Veilonellaceae and Lactobacillaceae are both from the Clostridium class (**Fig. 5a**). The Clostridia class is a major class in the early abdominal area stages and then decreases rapidly (**Fig. 4a**). On the other hand, Planococcaceae, family from the Bacilli class, rapidly increases in relative abundance in the Post-Bloat stages. Moraxellaceae also spikes in the Bloat stages (**Fig. 5a**), which coincides with the increase of Gammaproteobacteria (**Fig. 4a)**. In the anus, the early and Bloat stages have relatively similar composition. However, Gammaproteobacteria take over much of the composition in the Post-Bloat stages. These Gammaproteobacteria are almost exclusively from *Psychrobacter* and *Acinetobacter* genera. Both originate from the Moraxellaceae family, which also increases in the Post-Bloat stages (**Fig. 5b)**. In addition, Lactobacillaceae and Veilonellaceae generally decrease, while less abundant Corynebacteriaceae and Prevotellaceae display increases in the Bloat Two and Post-Bloat One stags respectively. The ear canal location has less relative abundance change at the class level, with Bacilli dominating in all stages in comparison to other locations. This is especially seen in stages before the bursting event (**Fig. 3)**. The Planococcaceae family, like the abdominal area, rapidly increases in the Post-Bloat stages (**Fig. 5c**), which may be the cause of these samples isolated from the rest of the ear canal samples (**Fig. 3)**. Staphylococcaceae spikes in the Fresh stage, while Veilonellaceae and Lactobacillaceae are most abundant at Placement (**Fig. 5c**). Finally, the nostril location showed a opposite trend with Clostridia and Bacilli in relation to the abdominal area, with a major increase in Clostridia in the Post-Bloat stages (**Fig. 4)**. Tissierellaceae, from the Clostridia class, is also seen to rapidly increase in the Post-Bloat stages (**Fig. 5d**). Interestingly, Clostridiaceae spikes in the Fresh stage, but then decreases and plateaus (**Fig. 6d**). Moraxellaceae has similar trend to the abdominal area sampling location, while Staphylococcaceae and Lactobacillaceae coincide with the ear canal (**Figure 6d**).

**Figure 5:**
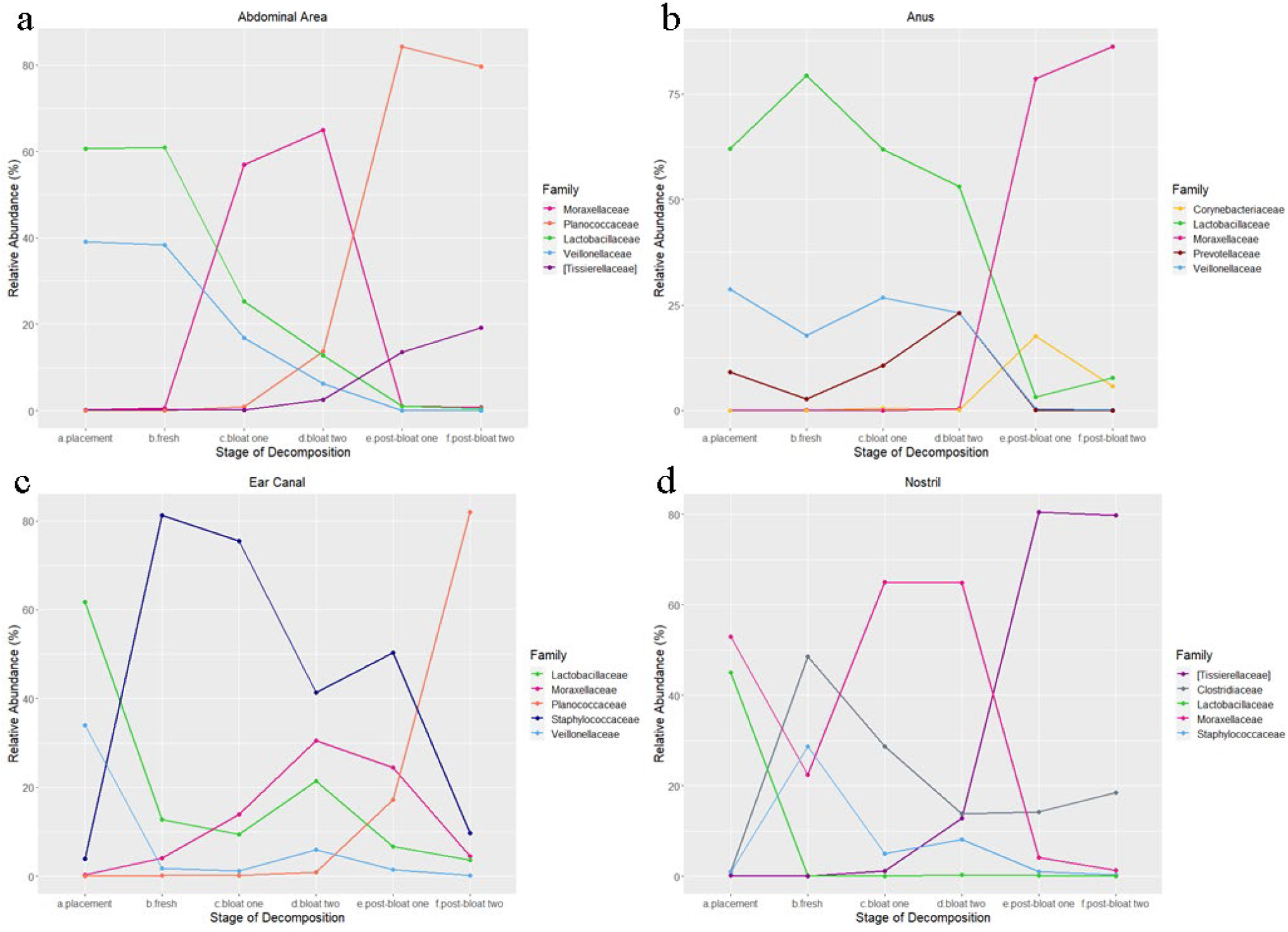
Top Five Prevalent Changing Family based on relative abundance (%) for **a**. Abdominal area, **b**. Anus, **c**. Ear Canal, and **d**. Nostril sampling locations. These line graphs were produced to better evaluate and isolate prevalent and shifting taxa.

**Figure 6:**
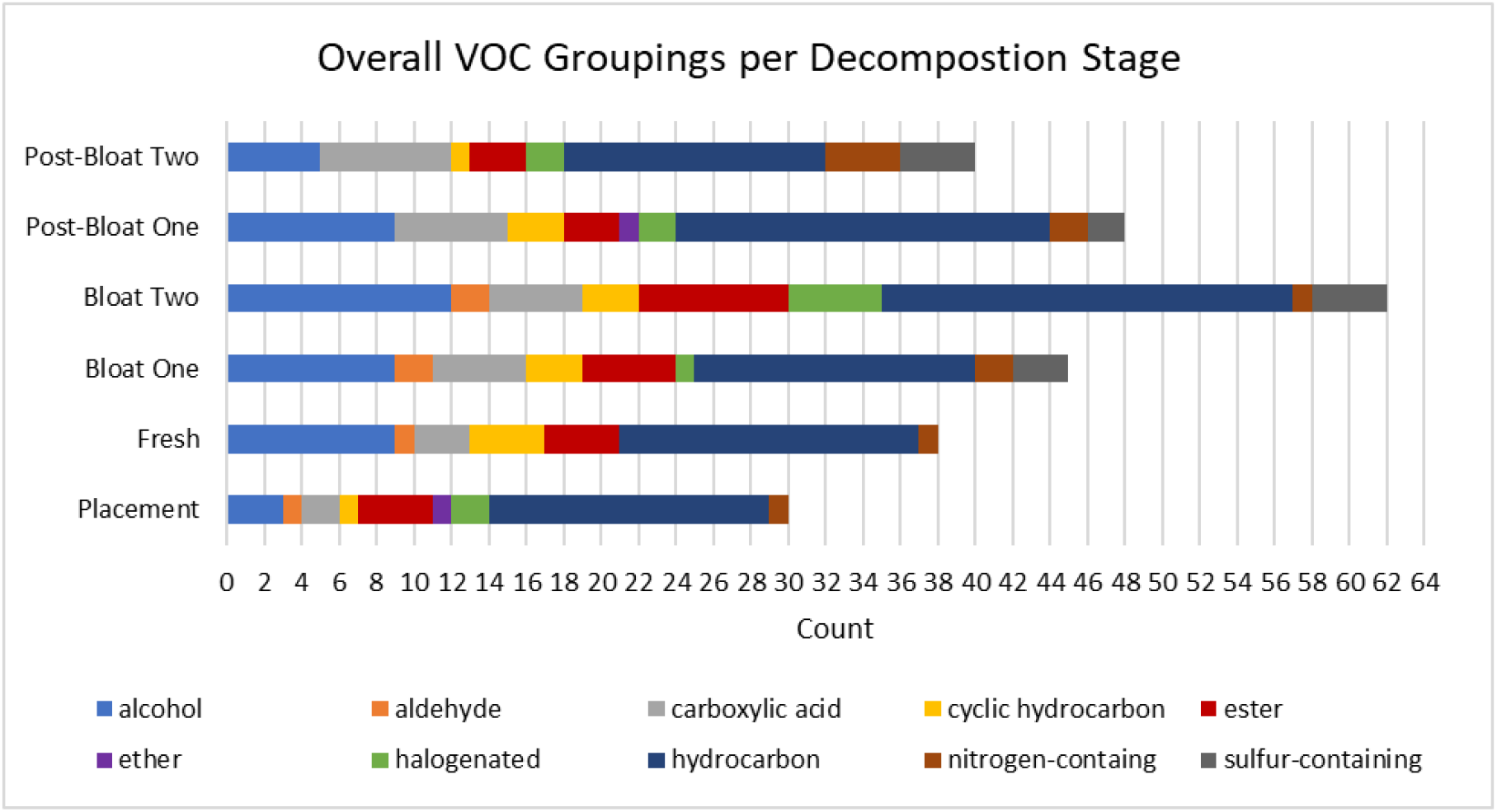
Stacked bar chart of functional group categories present in each stage of decomposition. The bar portions are measured in the number of VOCs categorized for each group, and functional groups were based on most defining functional group of VOC.

#### iv. Random forest analysis depicts the nostril sampling location as best predictor for stage of decomposition

In order to assess how accurately the stage of decomposition could be predicted using microbiome data alone, random forest analysis was performed. Stages of decomposition were grouped in various combinations by expanding or contracting decomposition phase parameters (**Table 1**). The different groupings were created to determine if using broader decomposition stage classifications would increase model accuracy. Out of bag (OOB) estimate of error rates was used to assess the accuracy of these different groupings in predicting stage of decomposition, along with comparing accuracy of the different sampling locations (**Table 2)**. OOB explains the amount of prediction error in the created model. In addition, predictor species for each Grouping and sampling location can be found in **File S3**. In general, despite the Grouping type, the anus swab location had the highest OOB estimate of error rates, while the nostril location had the lower estimates; the only exception was Grouping 4 with the ear canal having the highest estimate. Next, as the Groupings get broader (more stages condensed), the OOB estimate of error rate decreases. Broader groupings, like Groupings 3 or 4, showed a lower error rate than more specific groupings, like Grouping 1. However, the nostril and ear canal could be considered as fairly accurate predictors of decomposition stage with OOB estimates of less than 30%, regardless of Grouping type.

**Table 1:**
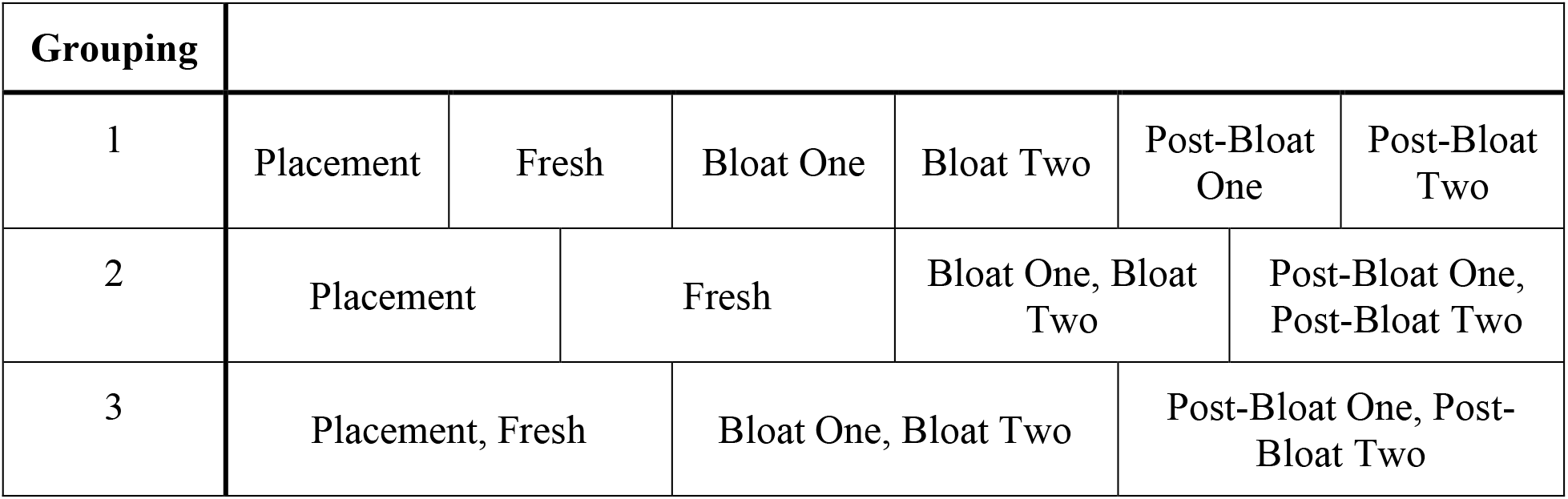
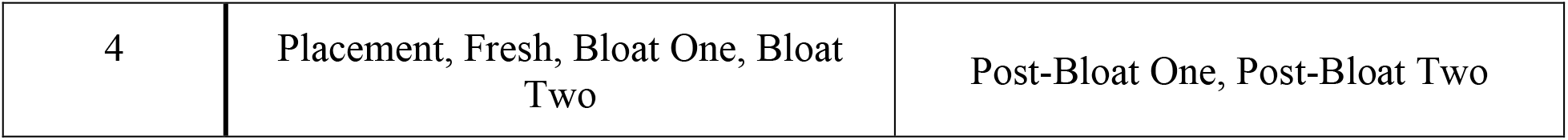
Recap of Groupings for random forest analysis (also see **Methods**).

**Table 2:** OOB Estimate of Error Rates of Different Random Forest Response Variable Groupings in Percentages (%). The smaller the OOB error rate estimates the more accurate the variable grouping is at predicting stage of decomposition based on the microbiome.

### B. VOCs

#### i. VOC functional groupings depict slight VOC profile change throughout decomposition

After removing VOCs found in the blank sample, there were between 30 and 62 identified VOCs per sampling day, peaking at the Bloat Two stage (**Fig. 6**). Most VOCs were classified as hydrocarbons. Either alcohols, esters, or carboxylic acids were the second most prevalent for each stage. Sulfur-containing compounds were observed after the fresh stage and into active decay. Aldehydes were seen up to the Bloat Two stage. Small peaks for cyclic hydrocarbons, halogenated compounds, and ethers are also present. Finally, nitrogen-containing compounds are present in all stages, but peak in the Post-Bloat Two stage.

#### ii. Temporal VOC patterns depicted using PCA plots

VOC profiles between stages of decomposition differed when plotted using VOC functional groups versus individual VOCs as loadings (**Fig. 7**). When functional groups were used as the loadings, decomposition stages were not as spatially separated (**Fig. 7b**), and VOC profiles exhibited only subtle temporal variations with decomposition stage (**Fig. 6**). In contrast, VOC profiles were more spatially distributed when using individual VOCs composition as the metric (**Fig. 7a**). The Bloat Two stage was isolated most likely due to the presence of increased and unique VOCs (**Table S1**). Additionally, Post-Bloat One is isolated, while Post-Bloat Two is slightly separated from the cluster containing Placement, Fresh, and Bloat One (**Fig. 7a**). Nitrogen-containing compounds are a driver of the Post-Bloat Two VOC profile (**Fig. 7b**), which is consistent with the increased number of nitrogen-containing compounds in comparison to the other stages (**Fig. 6**). Ethers are only found in the Placement and Post-Bloat Two (**Fig. 6**) and, therefore, their distributions trend towards ether and away from other stages (**Fig. 7b**). The other VOC functional groupings are indicative towards the Bloat Two stage due to the increased presence of VOCs, as mentioned previously.

**Figure 7:**
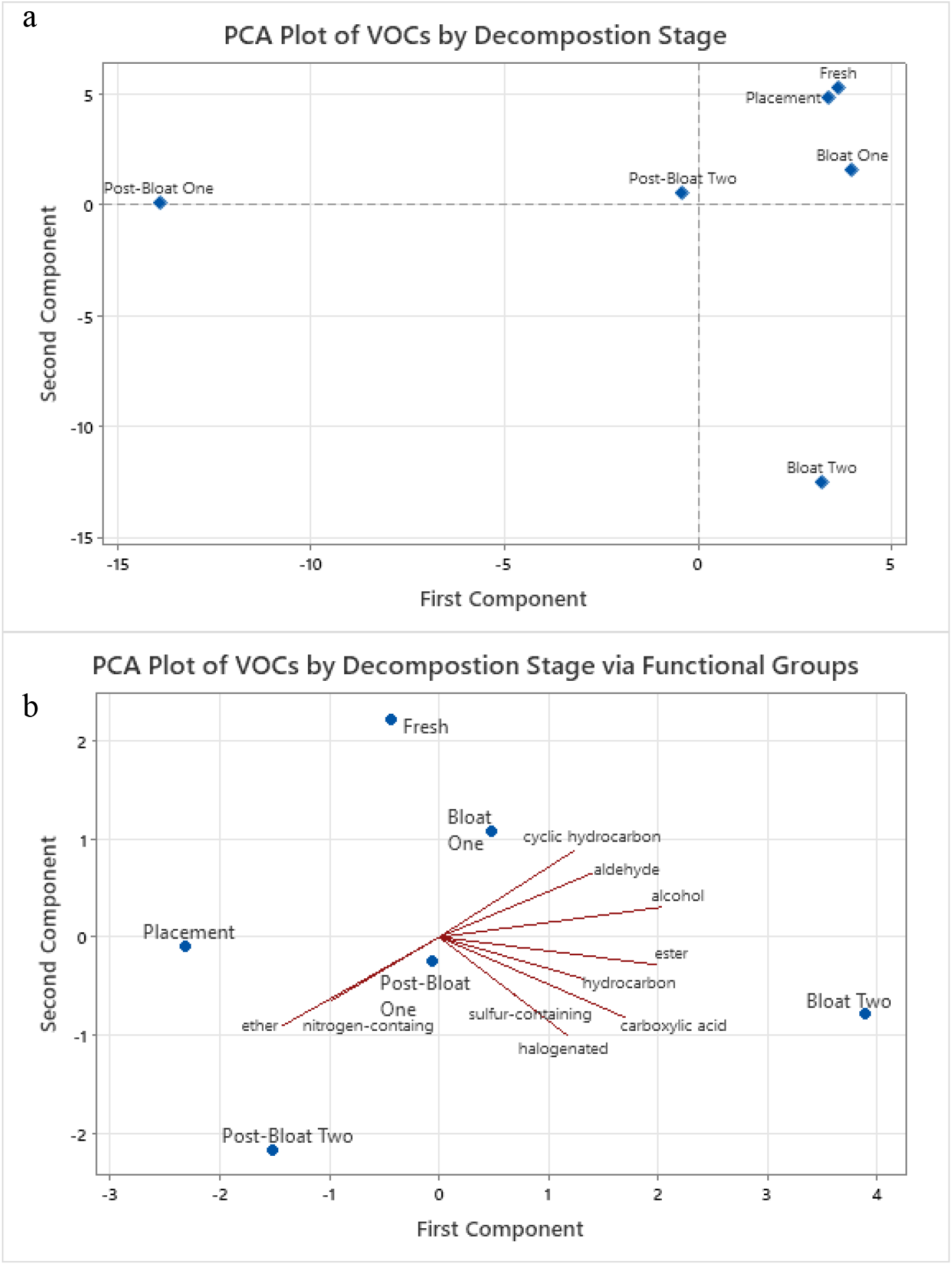
PCA Plots of Decomposition VOCs by Stage of Decomposition via **a**. Individual VOCs and **b**. Functional Groups. Loadings were calculated based on stage of decomposition. The PCA plots were created to determine the effect of specific VOC variability on stage of decomposition comparisons and if any category of compounds grouped towards stages of decomposition.

### C. Exploratory Linear Model Regression Analysis between Microbiome and VOCs

#### i. Selected VOCs and abdominal area top 30 bacteria families

The following VOCs were selected from the overall VOC profile for correlation testing to abdominal area bacteria families: nonanal, indole, dimethyl disulfide, dimethyl trisulfide, phenol, butanoic acid, and pentanoic acid. These compounds have been previously identified as characteristic to decomposition and were observed in more than one stage. From preliminary data analysis, the selected VOCs were categorized into one of three trend clusters (**Fig. 8a**). Unsurprisingly, VOCs in the same trend cluster showed higher correlation rates compared to those of in different trend clusters (**Figure S4**). Pentanoic acid and dimethyl disulfide are grouped together as Cluster 1 (**Fig. 8a**). Dimethyl disulfide was not observed in the Post-Bloat One stage, despite its reappearance in Post-Bloat Two. Both dimethyl disulfide and dimethyl trisulfide are common VOC markers and originate primarily from the breakdown of amino acid (6, 31). Along with dimethyl trisulfide, indole, butanoic acid, and phenol (Cluster 2) generally increase throughout decomposition (**Fig. 8b**). Phenol and indole, like sulfides, are known to attract insects and believed to result from the degradation of amino acids (4, 26). Butanoic acid, along with pentanoic acid, are carboxylic acids; these compounds are generally observed in the later stages of decomposition (25, 29). Carboxylic acids and other acids can be products of fermentation, a possible explanation to their later appearance (6). Finally, nonanal (Cluster 3) decreases throughout decomposition (**Fig. 8c**). It is a known insect attractant and has been identified in several decomposition studies with a range of hosts (28, 55, 56).

**Figure 8:**
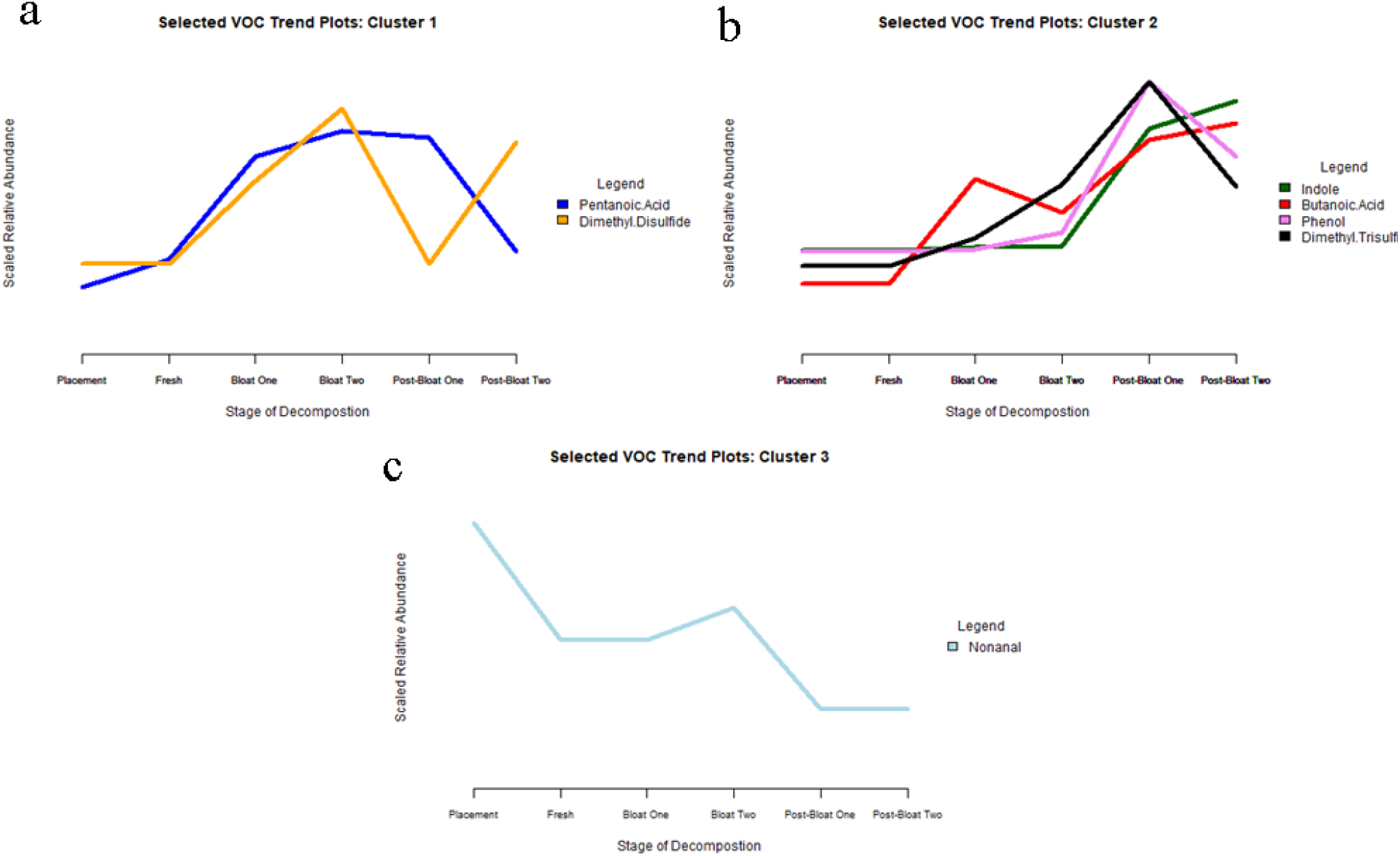
Line plots of selected VOCs based on scaled relative abundances. Selected VOCs are categorized into one of three clusters: **a**. Cluster 1 (Pentanoic Acid and Dimethyl Disulfide), **b**. Cluster 2 (Indole, Butanoic Acid, Phenol and Dimethyl Trisulfide), **c**. Cluster 3 (Nonanal).

The abdominal area location was selected because of the representative and exposed nature of the sampling location. On the other hand, the ear canal, nostril, and anus are small and more confined sampling locations. The abdominal area likely produces the majority of VOCs detected before, during, and after the bursting event. Additionally, after the bursting event, the displaced internal organs were also swabbed for this sampling location. The displacement of the internal organs may have introduced new bacteria that contribute to the overall VOC profile. The 30 most prevalent families from the abdominal area (**Table S4**) were selected to run linear regression models against the selected VOCs. In the same manner as the selected VOCs, correlations between bacteria families were assessed. A higher portion of pairwise correlations between bacteria families are positively correlated versus pairs showing no or negative correlations (**Figure S5**).

#### ii. Pairwise linear regression models produce exploratory results showing positive correlations between selected VOCs and abdominal area bacteria families

Pairwise linear regression models between selected VOCs and prevalent abdominal bacteria families were performed in order to assess any correlations, indicating possible associations between bacteria and VOCs throughout decomposition. The heatmap from **Fig. 9** shows that each selected VOC is positively correlated to more than one bacteria family, which is indicated by strong blue cells. Correlation patterns are closely tied to VOC trend clusters and groupings of bacteria families to produce visual correlation clusters (**Fig. 9**). Four major positively correlating clusters are present: 1) Nonanal positively correlates with the bacteria families Prevotellaceae through Veilonellaceae; 2) Dimethyl trisulfide has a strong positive correlation with Carnobacteriaceae through Planococcaceae; and 3) Dimethyl disulfide and 4) pentanoic acid correlate with two clusters of bacteria families of Peptostreptococcaceae though Corynebacteriaceae and Bacteroidaceae through Moraxellaceae. Due to opposing trends between VOC trend clusters (**Fig. 8**), strong negative correlations are often seen on other selected VOCs when strong positive correlations are depicted (**Fig. 9)**. For example, Prevotellaceae through Veilonellaceae have a strong positive correlation with nonanal, but a strong negative correlation with the other VOCs.

**Figure 9:**
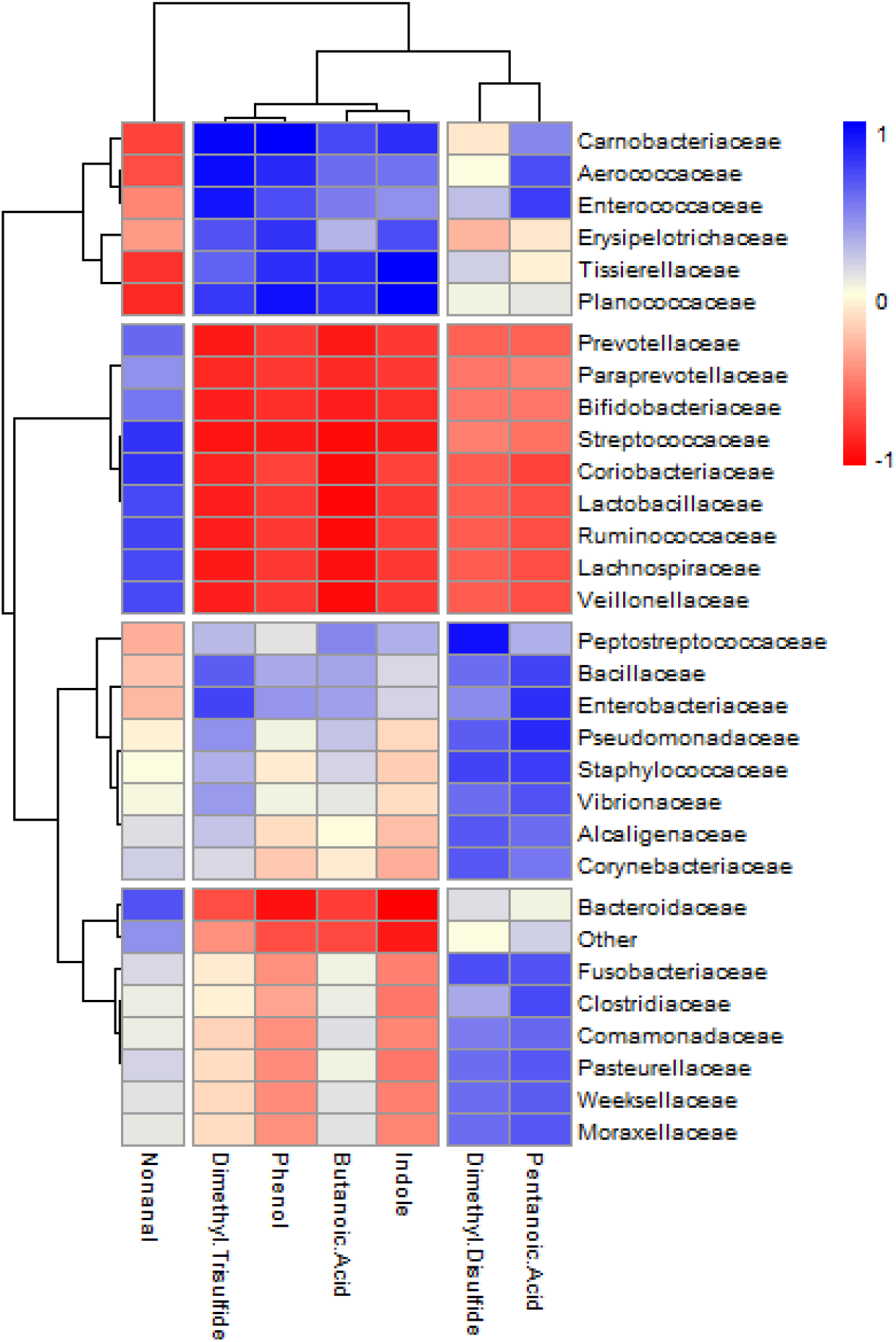
Heatmap depicting correlations, based on Pearson’s correlation, between pairs of bacteria families (rows) and selected VOCs (columns). Strength of correlation is shown through gradient of color scale: strong blue indicates strong positive correlation, strong red indicates strong negative correlation, and white indicates no correlation. Black bars on the left and top of the heatmaps are dendrograms, showing similarity of bacteria families and selected VOCs based on correlation patterns.

In addition to the visual representations of correlations between VOCs and bacteria genera, p-values (unadjusted and adjusted) were calculated to determine if the associations were significant or not (**Table S5**). Because of the limited number of sampling time points, this portion of the analysis is considered exploratory. Furthermore, constant peak area intensities like indole between Placement and Bloat Two peak can inflate significance of results. Finally, because only linear regression models were tested, possible non-linear trends or associations could not be explored. The families Planococcaceae and Tissierellaceae were positively correlated with indole (adjusted p-values < 0.001) (**Table S5**). The family Carnobacteriaceae was associated with both phenol (adjusted p-value < 0.001) and dimethyl trisulfide (adjusted p-value <0.01) (**Table S5**).

## IV. Discussion

Here, we identified how the microbiome and VOC profiles shifted during the course of indoor decomposition. We observed divergence in microbial communities for the different sampling locations, while also observing temporal shifts VOCs across decomposition timepoints. In addition, an exploratory linear regression analysis identified VOC-bacterial candidate pairs, illustrating their connection throughout decomposition.

Alpha diversity alone is not an informative metric to distinguish between decomposition stages. Based on past research, alpha diversity strongly depends on donor type, environment, and the duration of study (10, 12, 13, 34), which is consistent with trends from the indoor swine. Although most sampling locations produced significantly different results based on overall comparisons of alpha diversities between stages, pairwise stage comparisons produced few statistically significant comparisons, which could be due to the limited number of samples per location and per stage. In addition, inconsistent with our results, a prior study also identified that alpha diversity did not differ by sampling location (12).

With regard to microbiome compositional changes across locations and timepoints, it was not surprising to see Bacilli, Clostridia, and Gammaproteobacteria classes to be the main contributors. These three classes are a part of the Firmicutes and Proteobacteria phyla, both of which are repeatedly seen in decomposition studies (20, 57, 58). Despite the location, all Placement samples clustered together, which may be due to all sampling locations being epinecrotic, which includes the skin and superficial orifices (**Fig. 3**) (59). As decomposition progressed, communities shifted significantly depending on the sampling location, also seen in mice hosts and human cadavers (10, 12, 13). These differences may be possibility due to difference in nutrient sources, like oxygen, or bacterial interactions. Additionally, it can be supported that the indoor environment does not affect how the microbiome reacts during decomposition because temporal shifts are observed in our study, as with other microbiome decomposition studies (which are mostly in outdoor environments) (8, 10, 12-20, 58). However, due to the indoor environment, sampling only occurred up to the active decay stage because a significant amount of time is needed to reach the dry remains phase (1). Future work will need to address whether microbial communities continue to transform over a longer time period, as it has been observed taxa can be most unique in the dry remains stage (20).

First, the abdominal area shows a decrease in anaerobic bacteria (Clostridia class) with the onset of bloat, while the Moraxellaceae family increases, and then later the Planococcaceae family, which are more aerobic tolerant taxa. The Planococcaceae family, which dominates the Post-Bloat stages, is found in a variety of habitats (e.g., soil, water) (60). It is not a common decomposition related taxa; however, it was identified in a soil samples around a buried decomposing body (61). The abdominal area Post-Bloat samples included swabbing of the exposed internal organs displaced from the bursting event, leading to possible increased oxygen levels in the area swabbed. Furthermore, the sudden shift after the bursting event and increase in more aerobic tolerant bacteria may be due to this event (12).

Next, the anus sampling location differed from others because changes in community composition were minimal until after the bursting event. Although the anus location was not physically affected by the bursting event, the event is a significant decomposition landmark and is often associated with bacterial community shifts (10, 12, 20). The opportunistic pathogens, *Acinetobacter a*nd *Psychrobacter*, appearance in Post-Bloat stages may be due to the changing nutrient availability at this point in decomposition (8). *Psychrobacter* bacteria are psychrotolerant, while *Acinetobacter* are commonly found in soil environments (62-64). Moraxellaceae, the family of *Acinetobacter* and *Psychrobacter*, is a commonly identified family found throughout decomposition. In concordance with the family’s behavior in this study, the stage in which the family is identified in varies based on environment, sampling location, and host (10, 13, 20). Furthermore, the anus Post-Bloat samples show similarity to other sampling locations’ mid-study samples than other Post-Bloat samples, indicating the anus microbiome may be transform slower through decomposition.

Despite less apparent change at the class level, families like Planococcaceae and Staphylococcaceae, most likely drive the ear canal community shifts. Planococcaceae has been observed in other studies with an increase in later stages, as with other sampling locations in this study (13, 20). Staphylococcaceae bacteria, which peaks at the Fresh stage are facultative anaerobes, can use a wide variety of nutrient sources, and are common to mammalian skin microbiomes, making it unsurprising to discover it in the ear canal (65). Staphylococcaceae was also observed in a declining trend from a previous study in decomposing swine (34). Lactobacillaceae, prevalent in the Placement state, include lactic acid producing bacteria and may be a part of the inherent microbiome of the swine (66). However, Lactobacillaceae is usually associated with the gut microbiome, while it was found in all sampling locations in this study (66). Contradictorily to previous observations, Lactobacillaceae increased into the bloating phase of decomposition (10).

Finally, for the nostril canal, Tissierellaceae, from the Clostridia class, has been identified in decomposing bone samples, but little is known about the bacteria family (58, 67). Further exploration into the spike of Tissierellaceae, reveals that the family is primarily composed of the *Peptoniphilus* genus, an anaerobic dominate genus (68). Also, Clostridiaceae is a common family associated with decomposition, especially with other animal hosts (10, 13, 20, 34); its decline through the rest of the stages may be due to decreasing nutrient sources such as glucose (69). Lastly, Veilonellaceae, an obligate anaerobic family from Clostridia, has not been previously identified in other decomposition studies. Veilonellaceae is primarily found in earlier stages of all sampling locations, supporting that it may be a part of the swine’s inherit microbiome (70). Principally for the nostril sampling location, multiple shifts between anaerobic and aerobic dominate bacteria occur, unlike previous observations (12). More complex interactions or resource dependencies may be transpiring beyond oxygen availability producing these community changes.

Due to these microbiome shifts during decomposition, the microbiome can be used as a tool for determining PMI. Random forest analysis was used to evaluate this potential and found some sampling locations and stage groupings were more effective than others. Specifically, the nostril sampling location seemed to be the most predictive of decomposition stage, while the anus sampling location was the least accurate. The resolution of point in the decomposition process that could be predicted was limited, with higher errors observed when specific timepoints were predicted rather than broader timepoints (e.g., before and after the bursting event). Although current resolution is limited, these results demonstrate promise for the microbiome to be a useful marker for time since death. Because the process of decomposition is more accurately continuous rather than phase like (19), increasing sampling points and replicates would further improve these predications.

In reference to overall VOC production, the number of VOCs most likely spiked at peak bloating (Bloat Two) due to microbial activity being at its peak around this point. In addition, gas leakage is likely in the build up to the bursting event (8). Although hydrocarbons originate from the decomposition of lipids and are seen in most decomposition studies, they are not necessarily dominant nor persistent VOCs as in the current study (22, 23, 71-75). Another explanation in comment from another paper (76). It is unlikely the young swine in this study had significant adipose tissue reserves; therefore, the enclosed indoor environment may have facilitated capture of volatile hydrocarbons in contrast to other studies. The presence of sulfur-containing VOCs, main contributors to more complex odors associated with decomposition, during bloating and into active decay aligns with previous studies (4, 26, 29). Interestingly, aldehydes are more commonly seen in later stages in contrast to our observations (22, 25, 26, 29). Behind hydrocarbons, alcohols are a prevalent and common decomposition VOC category and can originate from a variety of macromolecule types (6, 22, 72, 77). Nitrogen-containing compounds include indole, a very common decomposition VOC, along with other nitrogen-contain compounds that are not observed in more than one stage. These different nitrogen-containing compounds may have not been consistently detected due to low detectability (27, 72). Ethers and carboxylic acids, derived from carbohydrates, are also commonly present in VOC decomposition profiles (22, 31).

The PCA plot (**Fig. 7**) supports that VOC profiles can change through decomposition. Temporal evolution through decomposition is supported by other studies like Stadler et. al. 2013, Forbes, et. al. 2014, Irish et. al. 2019, and others (23, 25, 27, 72, 75). VOC profiles through decomposition differ less based on VOC functional groupings versus individual VOC present. VOC abundances may be more reliable in correlating VOC profiles and decomposition stage, in part, because many individual VOCs can be categorized in more than one group, thus affecting VOC functional group trends.

Pairwise linear regression models were employed to probe potential associations between specific bacteria and VOCs. From this exploratory analysis, bacteria families that were identified to be positively correlated to selected VOCs may be potential producers, while bacteria families that had negative or weak correlation with selected VOCs may not. Planococcaceae and Tissierellaceae both were associated with indole significantly, and, unsurprisingly, they were strongly correlated to other VOCs of the same trend as indole. Both Tissierellaceae and Planococcaceae have been previously associated with indole (34, 78), and were identified are prevalent changing taxa from the abdominal area sampling location. Because these families were considered prevalent changing taxa from the abdominal area sampling location, these pairings may be promising contenders for future work regarding these bacteria-VOC relationships. Moreover, Planococcaceae may be a stronger contender as a producer of indole due its higher abundance levels, which can strength support of the association. Carnobacteriaceae had significant positive correlation with phenol and dimethyl trisulfide, but the family has not previously been identified to be associated with either VOC. Like indole, phenol does have constant measurements in early decomposition stages, which may have skewed p-values. Other prevalent changing families from the abdominal area like Lactobacillaceae and Veilonellaceae were positively correlated to nonanal. Again, this is predictable because these bacteria families and selected VOC all have general decreasing trends throughout decomposition. Moraxellaceae was strongly correlated to dimethyl disulfide and pentanoic acid, but these associations and the previously mentioned ones have not been observed before. It is important to note that these linear regression models only show association, not necessarily causation.

Again, due to the nature of the statistical analysis and data limitations, linear regression models are to be considered exploratory. Although many pairwise correlations between bacteria and VOCs were not statistically significant, heatmaps still revealed strong correlations between multiple bacteria family and VOC pairings. Both the microbiome and VOC profile showed trends through decomposition individually; the correlation analysis provide insight that these trends may be interconnected. Because a bacteria taxa and VOC can have strong positive correlation without a bacteria taxon to be known to produce a VOC, it is important to confirm these correlations with further research.

## V. Conclusion

A primary objective of the current study was to characterize the microbiome and the VOC profile of a decomposing body in an indoor environment. A shift in the microbiome was observed throughout decomposition, which supports the potential use of bacteria to determine the PMI. However, the observed shift is sample location dependent, and the nostril sampling location was determined to best at predicting stage of decomposition based on the microbiome. VOCs were concordant with other decomposition studies and individual VOC trends may help more with PMI determination. Pairwise linear regression models predict a significant correlation between the abdominal area bacteria and VOCs and suggests specific bacteria-VOC pairings may be important markers of decomposition stage or progress.

## VII. Acknowledgements

The authors express their appreciation for the ongoing support of Dr. Reena Roy. We would also like to thank the PSU Statistics Department, specifically Grant Hopkins, for his assistance on the statistical analysis. We are grateful to Megan Morris and Peter DeMartino for their support throughout the project. Finally, Dr. Craig Praul from the PSU Genomics Core Lab and Aswathy Sebastian from the PSU Bioinformatics and Genomics Consulting Center assisted and gave advice with sequencing and initial bioinformatics work respectively.

## X. Supplemental Materials

**Supplemental Table 1:** VOCs Identified in Each Stage of Decomposition.

**Supplemental Table 2:** Kruskal-Wallis Rank Sum and Dunn Multiple Comparison Results comparing Stages of Decomposition based on Shannon Diversity Index Alpha Diversities.

**Supplemental Table 3:** Kruskal-Wallis Rank Sum and Dunn Multiple Comparison Results comparing Sampling Location based on Shannon Diversity Index Alpha Diversities.

**Supplemental Table 4:** Top 30 Bacteria Families from Abdominal Area with Read Counts for Each Stage of Decomposition.

**Supplemental Table 5:** P-values from Linear Regression Models between Bacteria Families and Selected VOCs.

**Supplemental Figure 1:** Photograph of Indoor Enclosure with SMPE Fibers in place.

**Supplemental Figure 2:** Principal Coordinate Analysis Plot including Controls (Positive and Negative PCR and Extraction Controls and Experimental Air Controls).

**Supplemental Figure 3:** Shannon Alpha Diversity Measurements versus sample location for **a**. Placement, **b**. Fresh, **c**. Bloat One, **d**. Bloat Two, **e**. Post-Bloat One, and **f**. Post-Bloat Two. Points on boxes indicate specific sample measurements. Significant stage differences based on Dunn’s Multiple Comparison Test (p < 0.05) are denoted by “*.”

**Supplemental Figure 4:** Correlation Plot between Selected VOCs based on Pearson’s Correlation (shown by color scaled).

**Supplemental Figure 5:** Correlation Histogram between Bacteria Families based on Pearson’s Correlation.

